# “QuickStainer”: a rapid negative staining device for improved preservation of molecular structure

**DOI:** 10.1101/2025.10.11.681814

**Authors:** Vu Nguyen, Ruchi Gautam, Arun Kumar Somavarapu, Debabrata Dutta, Aditya Patra, Jinghua Ge, Christopher M. Yengo, Raúl Padrón, Roger Craig

## Abstract

Negative staining is a widely used technique for observing macromolecules and their assemblies by transmission electron microscopy. It is commonly employed to optimize specimens for cryo-EM. The stain, typically a uranyl salt, surrounds the structure, providing an outline view at about 20 Å resolution. Many macromolecules are relatively stable and rigid, and negative stain images provide a good representation of their structure. However, some are labile or flexible and their structure or assembly state is altered by binding to the carbon substrate on the grid before specimen staining. In these cases, the negatively stained appearance does not faithfully represent the structure in solution. This problem is reduced when samples are incubated on the carbon surface for short times (5 s) rather than typical times (30-60 s) before staining. To reduce disruption to a minimum, we have developed a rapid negative staining device (QuickStainer) using 3D-printed components, a stepper motor for precisely timed movements, and an Arduino-controlled interface to execute commands. QuickStainer produces consistent, reproducible results, achieving sample incubation times as low as 10 ms before staining. Tests show rapid adherence of molecules to the grid and greatly improved structural preservation of labile specimens compared with standard preparation protocols. The design of QuickStainer can accommodate inclusion of additional steps, such as timed incubation with enzyme substrate, before staining.

## INTRODUCTION

Negative staining is a preparative technique widely used to observe macromolecules by transmission electron microscopy (TEM) [1–7]. While cryo-EM has recently emerged as the method of choice for high-resolution structural analysis [8], negative staining retains an important role in optimizing specimens for cryo-EM and in providing rapid and inexpensive answers to questions that do not require high resolution. The technique typically involves applying a sample (protein complex or single molecule) in aqueous suspension to a carbon-coated grid, waiting for 30-60 s for its adherence to the carbon, then staining with a heavy metal salt solution [6, 7]. Uranyl salts (usually uranyl acetate or uranyl formate) are most frequently used and are known to “fix” specimens within 10 ms [9]. Excess stain is removed and the grid allowed to dry. Carbon-coated grids are frequently glow-discharged in a plasma before use to render the surface hydrophilic, aiding with spreading of the stain. Dried grids examined by TEM provide images of macromolecules surrounded by stain, revealing an outline of the structure at about 20 Å resolution. Stain typically binds more weakly to the protein than to the carbon substrate, leading to coining of the technique as “negative” staining.

Many proteins, complexes and assemblies (e.g. viruses) are relatively rigid and stable, and negative staining provides a good representation of their structure in solution. However, others, held together by weak interactions, are quite labile, and their structure is easily disturbed by binding to the carbon substrate, especially if it is highly charged through glow-discharge. Their appearance therefore may not faithfully represent the structure in solution (**Fig. 1a-c**). Examples in our laboratory are myosin molecules and filaments [4, 10]—highly flexible proteins and assemblies that are readily disrupted by binding to charged substrates [5, 11–15]. We are particularly interested in a motif formed by myosin II, in which myosin’s two heads fold back onto the tail and interact with each other, forming the interacting-heads motif (IHM, **Fig. 1e**) [16], an off-state found in myosin molecules of the cytoskeleton [15, 17–21] and in the thick filaments of relaxed muscle [22–26]. This “closed” conformation converts to an “open” structure, in which the heads no longer interact with each other or with the tail, when myosin is activated or when environmental conditions change (e.g. increased ionic strength; **Fig. 1e**) [27–29]. In our studies of the IHM in constructs of cardiac myosin II, we have encountered the problem that by negative-stain EM, few or no molecules show the closed conformation under solution conditions where most molecules are thought to be closed based on other solution techniques [10]: instead, most molecules appear open. Others have faced a similar problem, even with a more stable IHM (from smooth muscle myosin) [15]. There, the situation was improved by shortening the incubation time on the grid: this yielded a greater number of closed structures, suggesting that interaction with the grid surface was the problem [15]. Supporting this interpretation, we have found that reducing the time of incubation of our cardiac constructs from 30 to 5 s increases the number of closed structures, although most molecules still remain open [10]. Achieving low incubation times by hand is difficult and not reproducible, and times less than 5 s are not practicable. We have therefore designed and built a rapid, automated negative staining device, based on a computer-controlled rotary step motor, to bring incubation times below one second. Using this device (“QuickStainer”) makes short staining times possible (as low as 10 ms), greatly improving preservation of the closed IHM structure (**Fig. 1d**). The device is easily constructed and should be valuable for studies of diverse labile proteins and complexes.

**Figure 1.**
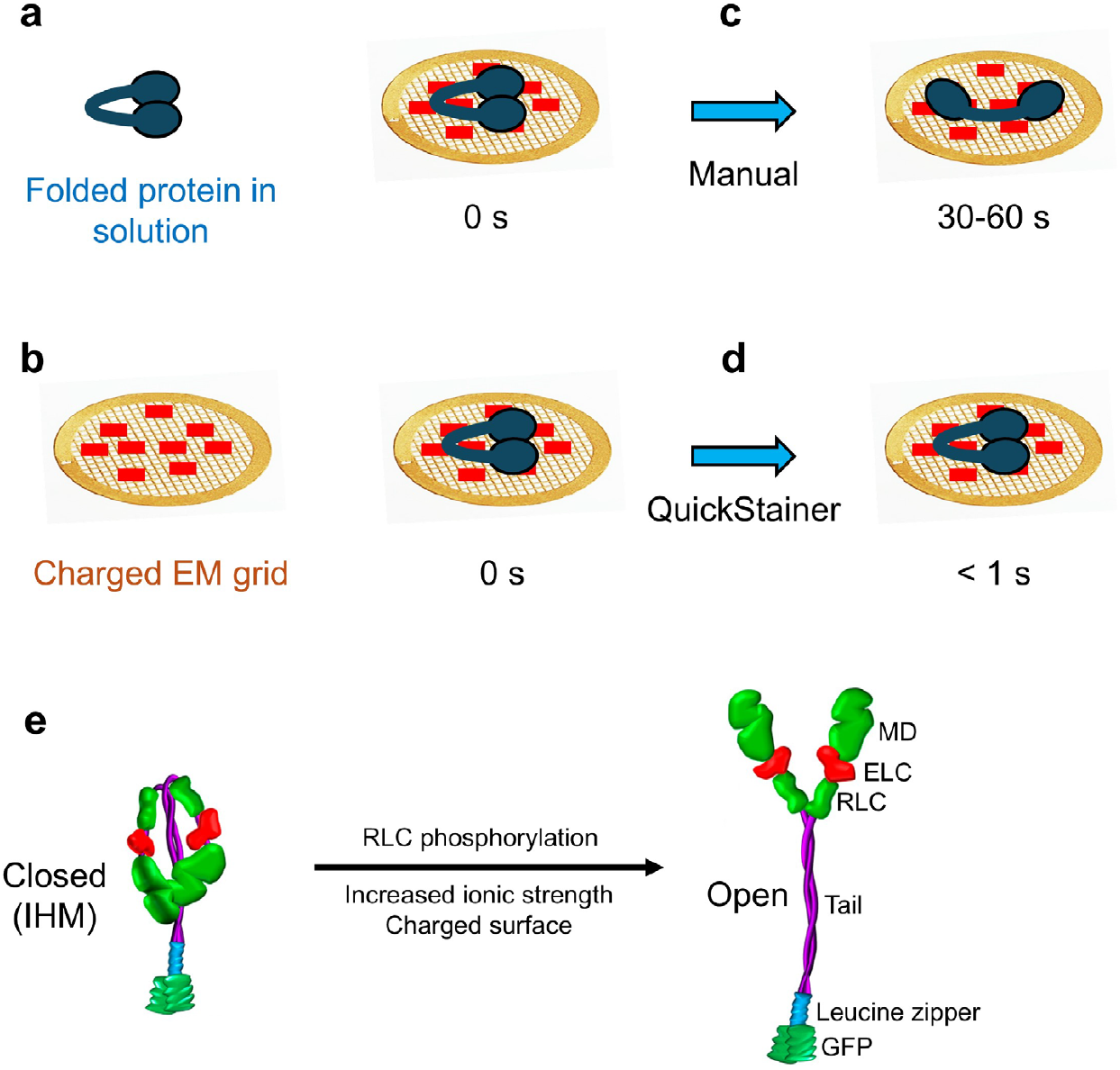
Labile proteins can change in structure upon binding to a charged grid surface. A protein that is folded in solution **(a)** binds to a charged grid surface **(b)**, causing it to change in structure if there is a long incubation time (30-60 s) before staining **(c)**. QuickStainer **(d)** minimizes incubation time (< 1 s), reducing change in structure. **(e)** Myosin II construct in closed form (off-state IHM) in solution converts to open form when activated (e.g. by phosphorylation), when exposed to high ionic strength (in vitro), or artifactually when it binds to a charged surface. MD, motor domain; ELC, RLC, essential and regulatory light chains; GFP, green fluorescent protein.

## METHODS

### Samples

Adeno-associated virus (AAV9873) was provided by the UMass Chan Medical School Electron Microscope Facility, apoferritin was obtained from Sigma-Aldrich (catalog #A3660), cardiac myosin constructs were prepared as described in [10], and smooth muscle myosin, provided by Dr. Mitsuo Ikebe, prepared as described in [30].

### Negative staining

Carbon-coated grids were obtained from EMS (CF-400-Cu-50). They were glow-discharged for 25 s at 15 mA in a PELCO easiGlo glow-discharger (Ted Pella) and used for staining immediately. Manual and quick negative staining were carried out with 1% w/v uranyl acetate in distilled water. For conventional (manual) negative staining, 5 µl of sample was applied to the grid for 5-30 s, followed by stain. The grid was then blotted dry by touching a filter paper to the edge of the grid, resulting in slow drying and a gradient of thick to thinner stain that allowed for optimal visualization of myosin constructs and other samples. The same drying procedure was used for quick-stained specimens. After complete drying, the grid was imaged on an FEI Tecnai Spirit Transmission Electron Microscope at 120 kV with a Rio 9 3K × 3K CCD camera (Gatan).

### Components of QuickStainer

The QuickStainer was constructed from 3D-printed parts made using an Original Prusa MK4S 3D printer (Prusa Research) and polylactic acid (PLA). The rotary stepper motor used to rotate the grid over the posts containing sample and stain was a Nema 17 Stepper Motor 2A 59Ncm, obtained from STEPPERONLINE. The electronics used to control the stepper motor was an Arduino Nano [A000005] microcontroller board and Stepper motor driver DM542T, obtained from Arduino and from STEPPERONLINE. The electronics were powered using LRS-150-24, obtained from Mean Well. Plans for 3D printing of the device parts are available on the Open Science Framework (see Data Availability).

### Assembly of QuickStainer

The two main components of the QuickStainer are the forceps holding the grid (**Fig. 2e, f**) and the stepper motor (**Fig. 2b**). Using 3D design (SolidWorks) and printing in PLA, a cylindrical mount was created to attach to the shaft of the stepper motor (**Fig. 2a, Videos 1, 2)**. This mount includes an insert designed to hold the forceps securely, fastened in place using a top-mounted screw. A custom platform was also designed and 3D-printed to support the sample and stain along the predetermined path of movement (**Figs. 2c, e, f, Videos 1, 2**). The platform features: 1. a central mount for securing the stepper motor; 2. radial posts positioned 18° apart to hold droplets of sample, buffer, and staining reagents; 3. a threaded center post for adjusting the height of the stepper motor (**Fig. 2c**). The 18° spacing between posts was based on the stepper motor’s 1.8° step angle (200 steps per revolution), enabling precise control by moving in 10-step increments. The stepper motor connected to a DM542T stepper motor driver, configured to increase the resolution from 200 steps per revolution to 800 microsteps per revolution through microstepping. The driver is powered by the Mean Well power supply and interfaces with the Arduino microcontroller (**Fig. 2d**).

**Figure 2.**
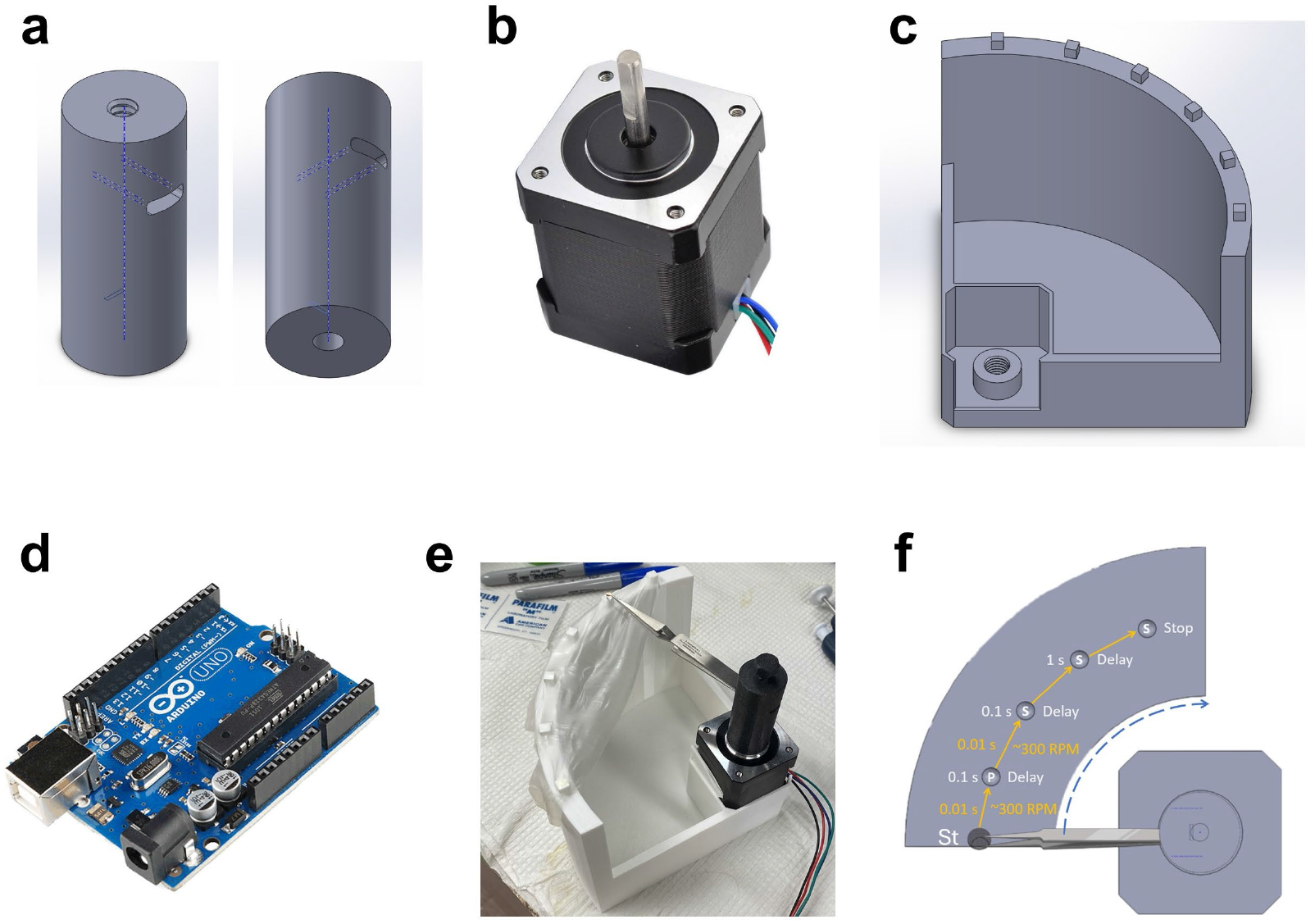
QuickStainer is constructed from four main parts. **(a)** a cylinder containing a slot for mounting the forceps that hold the grid, and a screw that inserts into the top hole to tighten the forceps in place. The cylinder attaches via the bottom hole seen in **(a)** to a stepper motor **(b)**, which fits in a square recess in the QuickStainer platform **(c).** The shaft of the motor is at the center of the circle defined by the posts, on which are mounted specimen and stain droplets. The threaded center post for adjusting the height of the motor fits into the threaded hole at the corner of the platform **(c)**. Rotation of the forceps is controlled by an Arduino board **(d)** attached to a laptop computer. A sheet of Parafilm is stretched over the posts of the platform, and 20 µL of sample and stain are applied to the posts **(e). (f)** The EM grid, held in forceps attached to the cylinder on the stepper motor, rotates from the starting position (St) to pick up protein specimen (P) on the first post before rapidly rotating to the second and later posts containing stain (S). Forceps are then removed from the device and the stain dried in the usual way. Delays can be programmed to vary incubation times at each post. See also **Videos 1, 2**.

We developed three Arduino programs to control the device: *Placement Program*, to align the forceps with the first post on the platform, the Starting post (**Fig. 2f**, St); *Reset Program*, to return the forceps to the starting position; *Experimental Program*, to execute the staining sequence by rotating the forceps through a series of four stations (**Fig. 2f, Videos 1, 2**), from the Starting post to: **1**. Protein Post – Rotates 18° (40 steps with 200-microsecond pulse periods) from the starting post to place the grid in contact with the protein sample. **2**. Stain Post 1 – Moves 18° to the staining position containing stain (uranyl acetate), which rapidly fixes [9] and stains the protein on the grid. A 100-millisecond delay ensures sufficient interaction time for initial staining. **3**. Stain Post 2 – Rotates another 18° (40 steps with 1000-microsecond pulse periods) to a second staining station, also containing uranyl acetate. Since fixation stabilizes the sample at Step 2, a longer, 1-second dwell time is used to enhance reagent exchange. The second stain post ensures removal of any protein components that may have detached from the grid and contaminated the stain at the first stain post. **4**. Stain Post 3 – Advances 18° to a third uranyl acetate station, where the grid remains until the forceps are removed, and the grid is blotted and dried as described for manual staining, above. The Reset Program is then initiated ready for the next run (**Video 2**). Note that Stain Post 1 could contain rinse buffer instead of stain, if desired, to avoid mixing of loose protein in the sample droplet with the first drop of stain. However, our goal was to transfer the grid to stain as soon as possible after picking up sample from the Protein Post to minimize structural change due to interaction with the grid surface. We therefore dispensed with any rinse step, which seems superfluous given the short duration in stain at step 2.

### Using QuickStainer

The following is our standard protocol for quick staining (**Videos 1, 2**). **1**. Apply parafilm to posts, providing a clean, hydrophobic surface for supporting the droplets without spreading. **2**. Program parameters for desired incubation time and speed and perform test run. **3**. Apply 20 µL water droplets to posts to simulate sample and stain droplets. **4**. Adjust height of center post so that grid will sweep across top of droplets. **5**. Perform test run. **6**. Pick up glow-discharged grid with forceps and attach forceps to cylindrical holder. **7**. Apply 20 µL sample and 20 µL stain to designated posts. **8**. Run Placement program to align grid to first post. **9**. Run Experimental program to stain grid. **10**. Run Reset program to return grid to initial position, near starting post, for easy removal of forceps. **11**. Remove forceps holding sample grid, blot grid to remove excess stain, and allow to dry.

### Image classification and averaging

Particles in images of myosin constructs and of folded, full-length (10S) myosin were classified automatically using the BlobPicker module of CryoSPARC v4.4, and class averages were calculated. The number of particles in different class averages was taken to indicate the frequency of each structure. Class averages of IHMs were considered to be open when there was no obvious interaction between the motor domains of the two heads and closed when the motor domains interacted and were folded back onto the tail **(Fig. 1e)**.

## RESULTS

### Protein particles bind to grid even with short incubation times

The QuickStainer works on the principle that limiting the incubation time of the sample on the glow-discharged carbon surface should reduce surface-induced disruption of specimen structure. Once the sample encounters the uranyl salt used for negative staining (usually uranyl acetate or uranyl formate), it is fixed within 10 ms and no longer changes in structure [9]. Typical incubation times with manual staining are 30-60 s [6, 7]. With short times, we might expect less sample to adhere to the grid surface. We therefore tested two samples (AAV and apoferritin) to determine how sample load was affected by incubation time.

For manual staining, AAV and apoferritin were applied to the grid for 5 s at concentrations that yielded a useful distribution of particles, with good contrast and an even spread of stain **(Fig. 3a, c)**. These served as controls. The same specimens were also stained with the QuickStainer, with shorter incubation times before staining. With 1 and 0.5 s incubation, particle distribution, concentration and stain spreading were similar to those for 5 s manual staining **(Fig. 3b, d)** after a moderate (2-3 fold) increase in concentration when necessary. Thus, the 30-60 s incubation times typically used for negative staining are not required: good sample load, distribution and staining can be obtained in ≤1 s.

**Figure 3.**
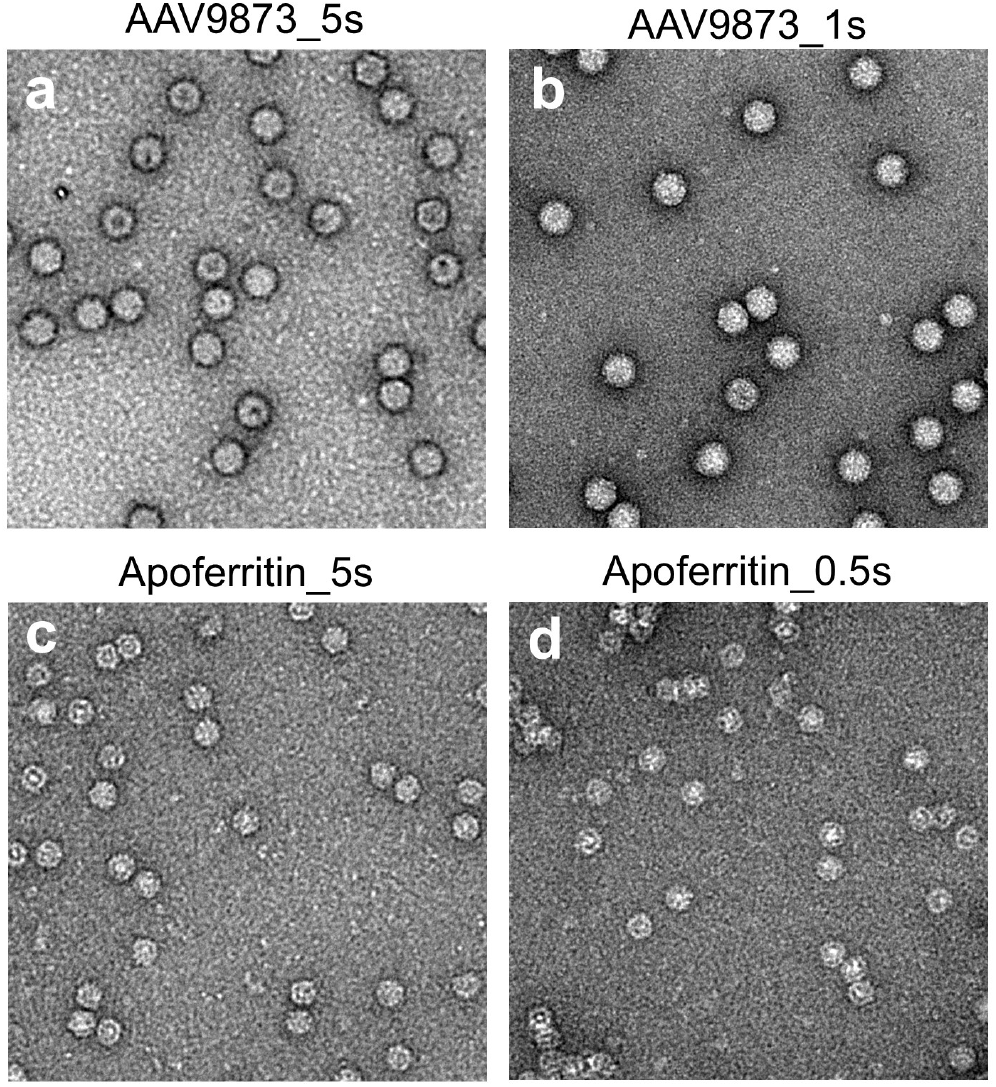
Protein particles bind to grid even with short incubation times. Because QuickStainer greatly reduces specimen incubation time on the grid compared with conventional staining, we checked whether a useful particle load could still be obtained with short incubations. **(a, b)** AAV applied for **(a)** 5 s (manual staining) and **(b)** 1 s (QuickStaining); **(c, d)** Apoferritin applied for **(c)** 5s (manual) and **(d)** 0.5 s (Quick). A useful load and particle distribution was obtained in each case for QuickStainer, similar to manual staining, and particles looked similar. When particle loads were reduced with short incubations, as occurred on some occasions, this could be corrected by a moderate increase in concentration (2-3 x).

### QuickStainer improves preservation of the closed IHM structure

We next examined our 2-headed construct of β-cardiac myosin II **(Fig. 1e)** under solution conditions (low salt) where FRET and ATP-turnover measurements suggest that > 90% of particles have a closed conformation [10]. Even when incubated on the EM grid for only 5 s before manual staining, most particles were in the open conformation (**Fig. 4a**, blue circles), while closed conformations were rare (pink circles). We compared these findings with shorter incubation times (0.5, 0.1, 0.01 s) using QuickStainer. This required an increased protein concentration to produce a useful load and distribution of protein (up to 1.5 µM for 0.01 s, compared with 100 nM for 5 s staining). With these shorter times, compact (closed) particles became common **(Fig. 4b-d)**. To quantify open and closed conformations objectively, we performed image classification and class averaging (CryoSPARC v4.4, **Methods**). The results supported our conclusions from the raw images. Closed structures were essentially absent with manual staining, but increased to almost 60% with 0.01 s incubation time **(Figs. 5, 6)**. This suggests that the closed structure is minimally disrupted with brief incubation, and can approach the numbers actually thought to exist in solution.

**Figure 4.**
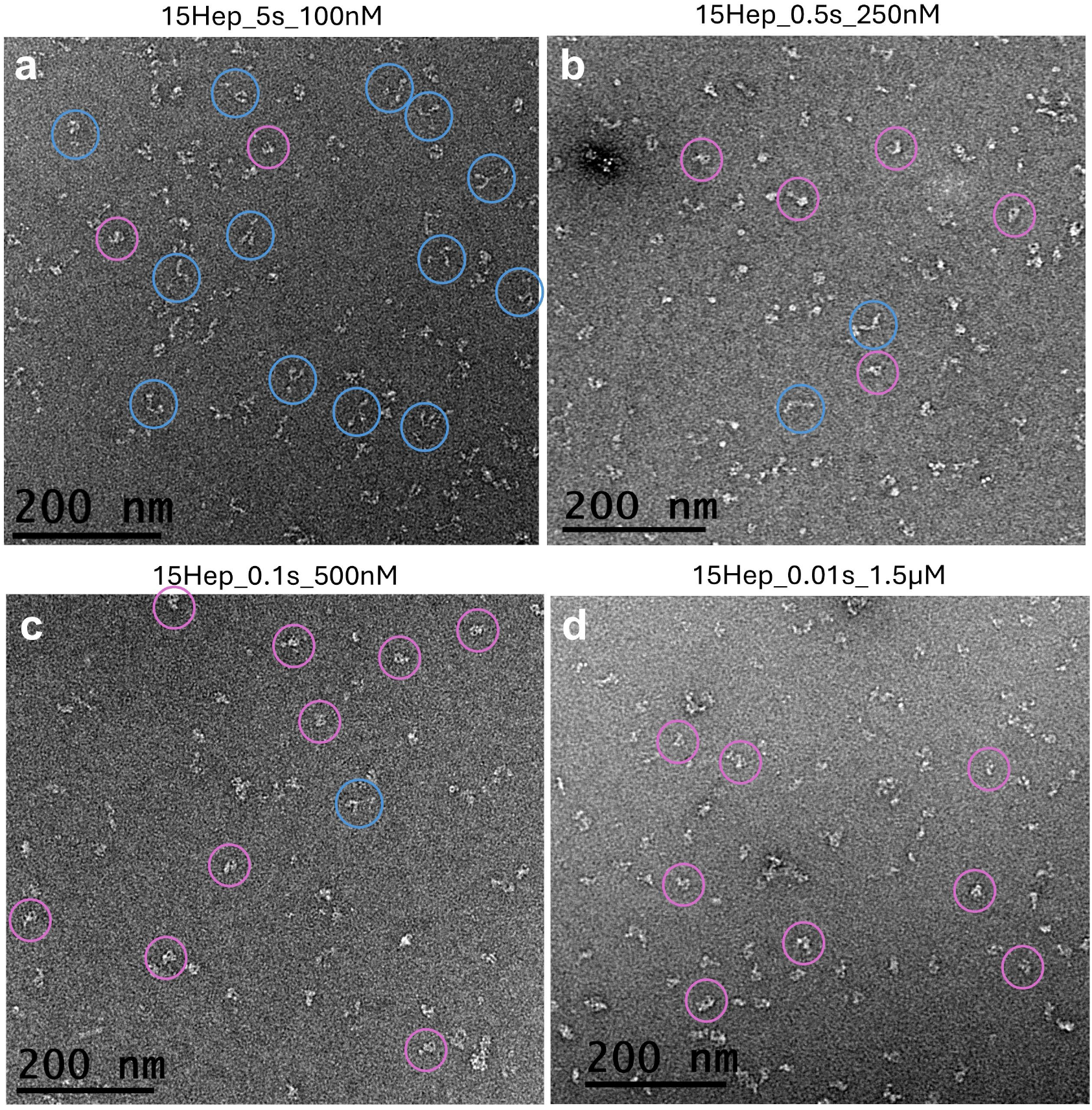
Reduced incubation time of myosin II construct produces more closed structures. A myosin II construct with 15 heptads of tail (15hep; **Fig. 1e**) was applied to a grid for **(a)** 5s (manual, 100 nM), **(b)** 0.5 s (250 nM), **(c)** 0.1 s (500 nM), and **(d)** 0.01 s (1.5 µM) before staining with uranyl acetate; **(b-d)** all using QuickStainer. There were more compact structures (pink circles) and fewer open (blue circles) as the incubation time decreased. The increase in protein concentration with shorter incubation times was needed to keep the final load constant. See also **Figs. 5, 6**.

**Figure 5.**
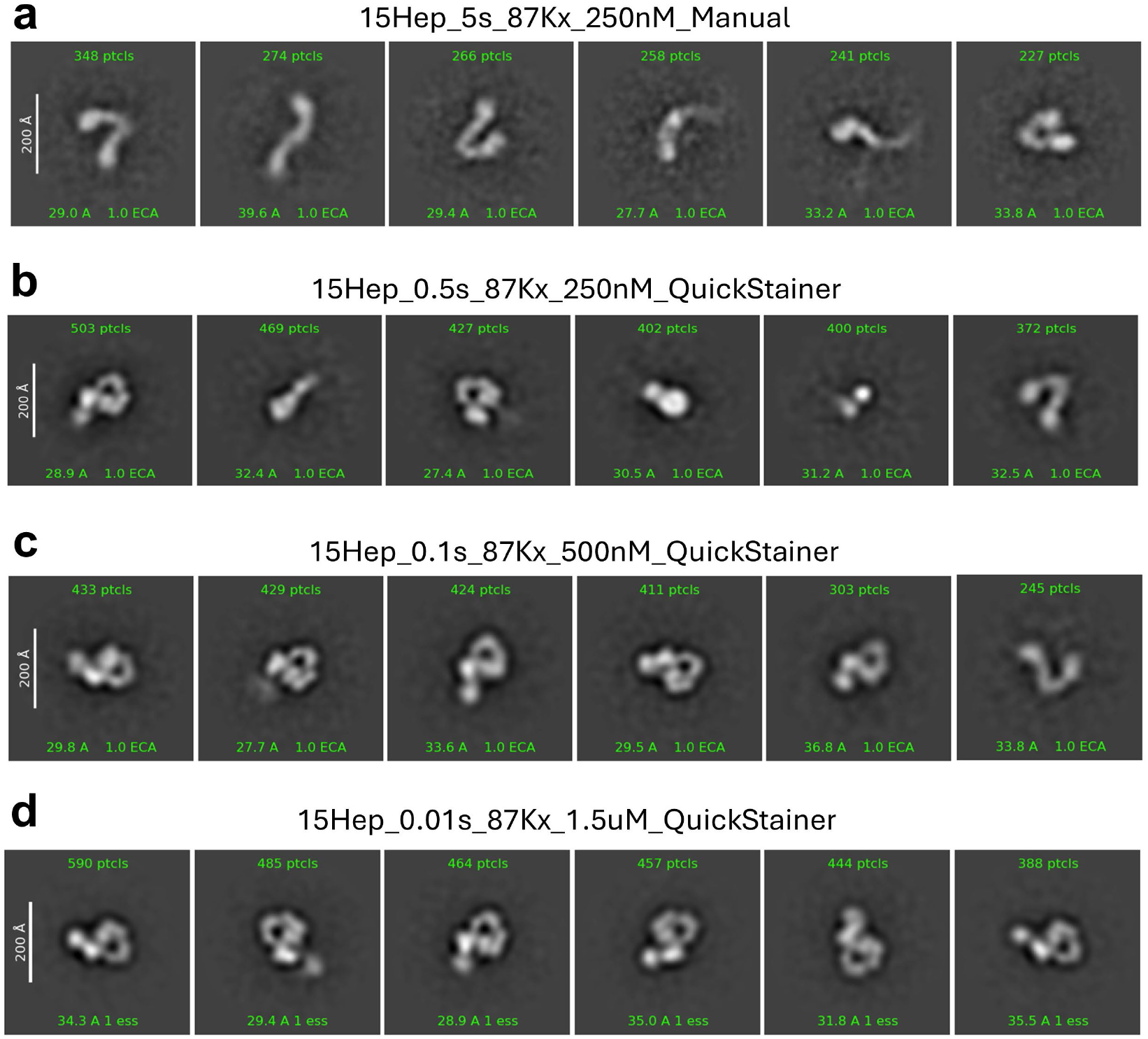
Closed and open conformations can be quantified by class averaging. Particles in images like those in **Fig. 4** were classified automatically using Blob Picker followed by Template Picker in CryoSPARC v4.4, and class averages calculated. The six most frequent classes are shown for 5 s, 0.5 s, 0.1 s and 0.01 s incubation times (number of particles at top of each box). With 5 s, all six averages are open structures; closed structures become more frequent with shorter incubation times, until the six most frequent classes are all closed with 0.01 s incubation.See **Fig. 6** for numerical comparison.

**Figure 6.**
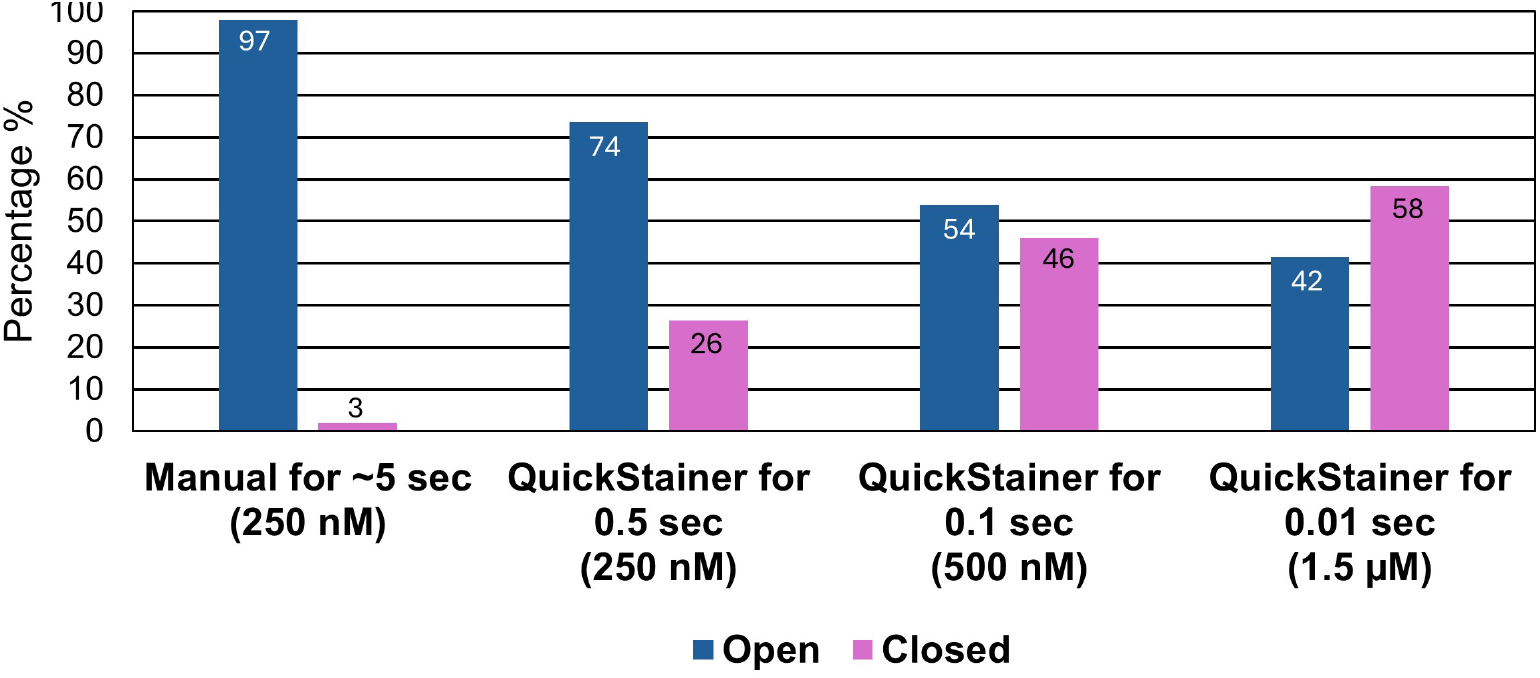
Increase in closed conformation with decreased incubation time. The graph shows the percentage of particles contributing to particle averages that have a clear open or closed conformation, using the top 6 (most frequent) classes (**Fig. 5**) and additional, less common classes (not shown). The number of closed structures increases with shorter times of specimen contact with the grid before staining.

### QuickStainer preserves the compact structure of full-length (10S) myosin II

To further evaluate the efficacy of QuickStainer, we carried out experiments on a different molecule. Full-length smooth muscle and nonmuscle myosin IIs fold into a compact structure in the off-state— referred to as 10S myosin, based on its sedimentation coefficient [27, 31, 32]. The heads fold back on the tail, in the IHM conformation, and in addition the 155 nm-long tail also folds, at hinges 1/3 and 2/3 along its length, wrapping around the heads and forming a compact association of the three tail segments [33–36], which fully shuts down activity [37]. When 10S myosin is examined by conventional (manual) negative staining on glow-discharged grids, it often shows a looped structure in which the tail segments are separate from each other, rather than the compact conformation where they interact; the heads may also lack head-head interactions [38]. A similar appearance is seen with rotary shadowing, where the molecules are supported on a charged mica surface before shadowing [27, 32, 39]. Strikingly, if the molecules are first crosslinked, in solution, before staining or shadowing, they show a fully compact structure [28, 33, 38]. These observations again suggest that weak interactions, here maintaining the 10S conformation, are disrupted due to binding to the charged substrate—thus providing a misleading idea of the structure present in solution. We tested whether QuickStainer overcomes this problem. When 10S molecules were applied to a glow-discharged carbon-coated grid for ~ 5 s before staining, most molecules (~70%) showed a less compact structure, usually without interactions between the heads **(Figs. 7a, c)**. With 0.01 s incubation time using QuickStainer, most molecules (~70%) now showed a compact, fully 10S structure (**Fig. 7b, d**), with heads folded back and tail segments together, similar to crosslinked molecules, confirming the importance of short incubation times for the preservation of labile structures.

**Figure 7.**
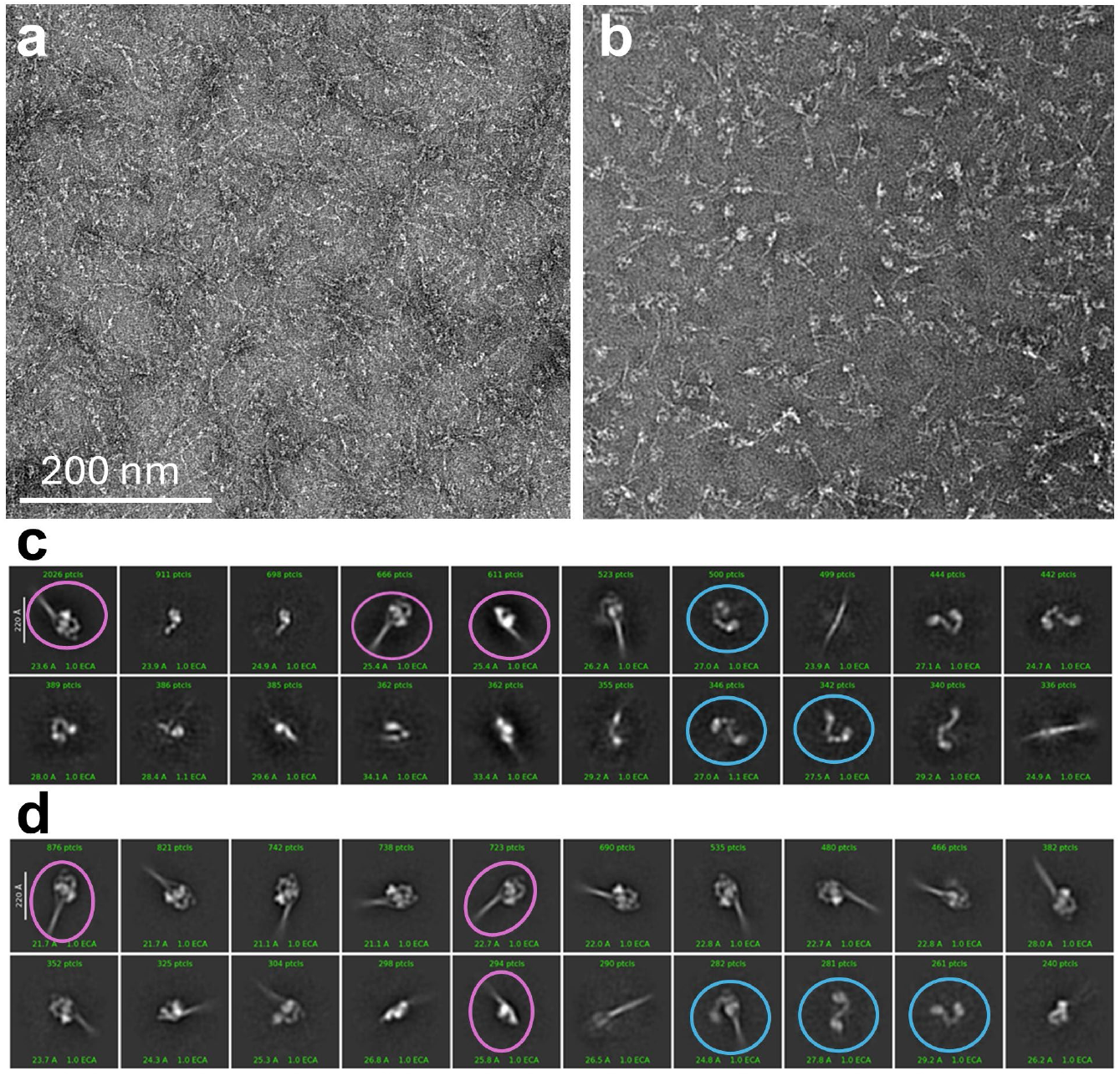
Testing QuickStainer with folded, full-length (10S) myosin. **a, b**. Smooth muscle myosin in the switched-off (10S) state negatively stained with **(a)** manual (5 s) and **(b)** QuickStainer (0.01 s) incubation times. **c, d**. Class averages from conditions corresponding to **a, b**, respectively. Only 3 class averages in **c** show definitive 10S features (pink) (see [33] for interpretation of images); the rest show single heads, double heads (non-interacting; blue) or are not identifiable. Sixteen class averages (all but the last 4) show 10S features in **d**. Overall, approximately 30% showed 10S features with 5 s and 70% with 0.01 s staining. The greater frequency with QuickStainer shows that short incubation times help to minimize structural disruption caused by binding of specimen to the charged carbon substrate. Note: in contrast to [33], the myosin was not crosslinked in these experiments.

### QuickStainer preserves the helical arrangement of myosin heads in thick filaments

In relaxing conditions, the thick, myosin-containing filaments from striated muscle have a helically ordered arrangement of heads in the IHM configuration [22, 40–43]. When studied by negative staining on glow-discharged carbon or rotary shadowing on mica, this delicate and labile helical organization is frequently lost, again apparently due to the charged specimen substrate used to support the sample, which is thought to disrupt the weak electrostatic interactions forming the IHMs and holding them in their helical positions [4, 11, 13]. We tested whether QuickStainer could reduce this problem in filaments, as it does in single molecules. When tarantula myosin filaments [44] were observed by manual staining (30s) on a glow discharged grid, heads were mostly disordered (90% of filaments) **(Fig. 8a, b)**. With short (1.0 s) incubation using QuickStainer, helical order was well preserved (~70% of filaments) **(Fig. 8c, d)**. Thus filamentous assemblies, like single molecules, adhere quickly to the carbon surface, and their delicate, flexibly attached heads are preserved in their known positions in solution if contact time with the grid surface is minimized.

**Figure 8.**
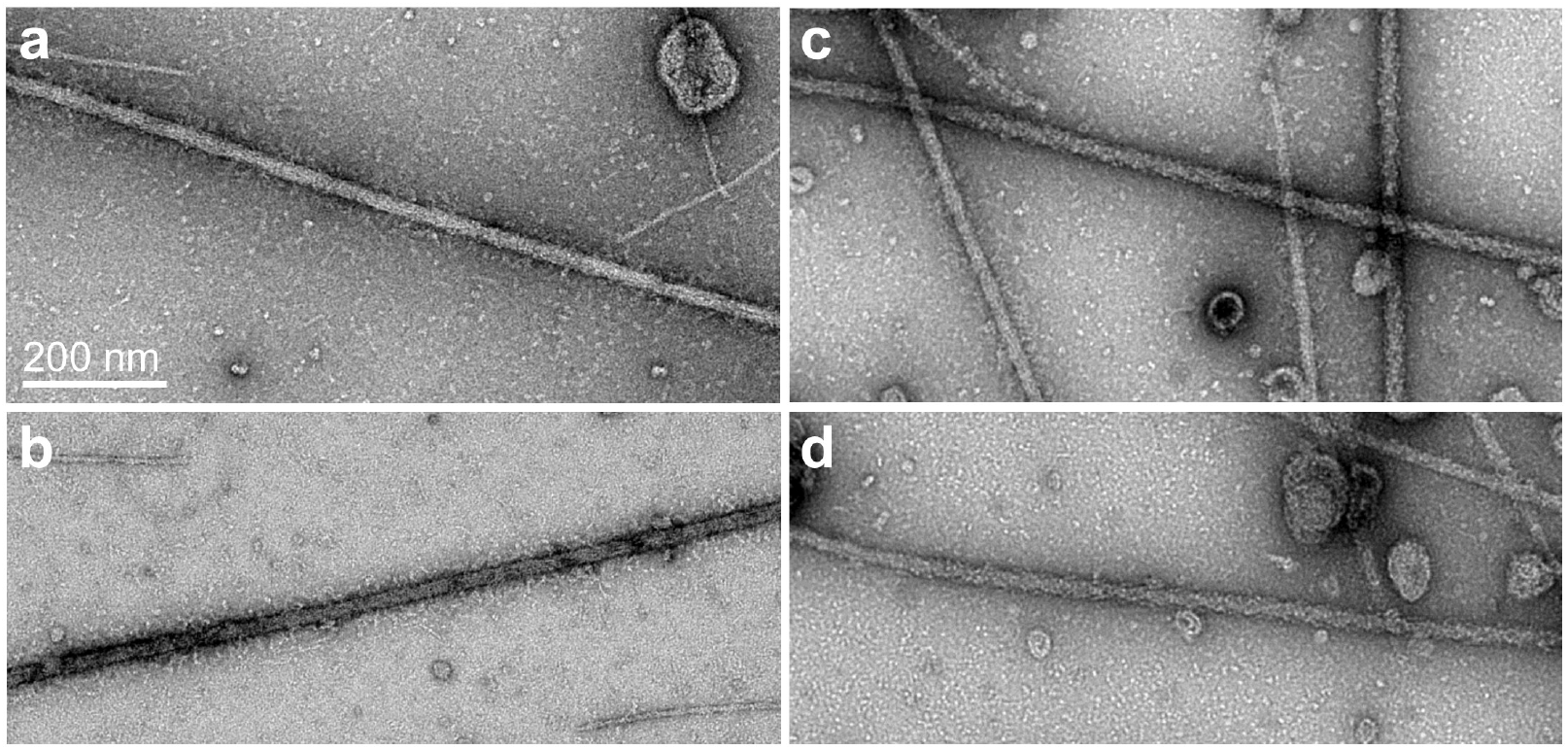
Testing QuickStainer with myosin filaments. **a, b**. Tarantula native myosin filaments incubated on glow-discharged grid for 30 s before manual negative staining with 1% uranyl acetate. **c, d**. Same as **a, b**, but with incubation time of 1 s using QuickStainer. In **a, b**, myosin heads form a fringe along the edges of the filament backbone (~90% of filaments), whereas in **c, d**, most filaments (70%) are helically ordered (myosin heads do not form a fringe, but are regularly arranged along the filament backbone forming a diamond-shaped pattern). All micrographs at same magnification.

## DISCUSSION

Labile proteins and macromolecular assemblies are easily disrupted when applied to charged EM grids used for negative staining, leading to incorrect interpretation of their structure in solution; an analogous problem occurs with shadowing, where molecules adhere to a charged mica surface, and in cryo-EM, when particles denature at the air-water interface [45]. We have shown that QuickStainer greatly reduces this problem, by minimizing the contact time of the sample with the grid surface before it becomes rapidly fixed (within 10 ms) by the uranyl stain [9]. The quick staining technique works for both molecules and molecular assemblies. We expected that short incubation times would cause a decrease in protein loads on the grid. Surprisingly, AAV and adenovirus particles adhered within 1.0 s in numbers similar to those found with conventional negative staining (5-30 s; **Fig. 3**), with only minor adjustments to protein concentration when necessary. In the case of our myosin construct, higher concentrations were required at the shortest incubation times (**Fig. 4**), which could be a limitation with QuickStainer if protein is in short supply. On the other hand, one could take advantage of this fact when imaging a weakly associated protein complex, which may tend to dissociate into its subunits at the low concentrations typically used for conventional negative staining. Dissociation can be reduced using higher concentrations, but this may produce an overloaded grid with standard (manual) negative staining procedures. Using the short incubation times possible with QuickStainer, high protein concentrations could be used, maintaining association of subunits, without overloading the grid.

### Pros and Cons of QuickStainer

QuickStainer has advantages in addition to minimizing disruption of protein structure caused by the charged carbon surface. In some of our observations of the IHM by conventional negative staining, we see both edge- and face-views of the structure, but the face-views typically have the same orientation—i.e. the molecule adheres to the grid with the same face (Fig. 6 – figure supplement 1 of [10]). A similar tendency was observed with the heads of extended myosin molecules observed by conventional negative staining [14], and preferred orientation is also encountered with cryo-EM [45]. With QuickStainer, we see face-views that are mirror images of each other (**Fig. 5d**), implying that both faces can adhere to the carbon substrate, an advantage that could be exploited for single particle 3D reconstruction. There are two possible explanations for the difference between fast and slow staining. Particles may be more stable on the substrate when adhering by the favored face; those adhering by the opposite face may denature during the prolonged incubation of manual staining and therefore not be observed. Alternatively, particles may bind only weakly by the opposite face, and dissociate and reattach by the favored side during incubation. Both scenarios are consistent with the improvements seen with short incubation.

Interestingly, the ability of quick-staining to minimize molecular disruption may sometimes mask useful insights into protein structure/properties that are obtained with slow staining. An example is our investigation of the dilated cardiomyopathy (DCM) E525K missense variant in the myosin head [10]. Our FRET observations suggest that the IHM is stabilized by this charge-reversal, through a strengthening of the electrostatic interaction between one of the folded-back heads and a negatively charged patch on the tail (**Fig. 1e**). Conventional negative stain EM is qualitatively consistent with this finding: while wild type (WT) shows few closed structures (IHMs)—similar to **Fig. 4a** here—this number increases with the variant [10] (**Fig. 9a, b, e**). In striking contrast, QuickStainer (0.01 sec incubation) shows a high number (~60%) of closed structures for WT (**Figs. 4d, 5d, 6**), for reasons we have discussed above, but little increase above this with the variant (**Fig. 9c, d, e**). Thus, slow (conventional) staining reveals the greater stability of E525K, because it (artifactually) disrupts the weaker WT structure, while the mutant is strong enough to partially resist the disruption. With QuickStainer, WT as well as E525K are both stable enough to remain in the closed conformation when contact with the charged substrate is short, hiding the difference in stability. In such situations, a comparison of manual and quick-stain methods is the most revealing.

**Figure 9.**
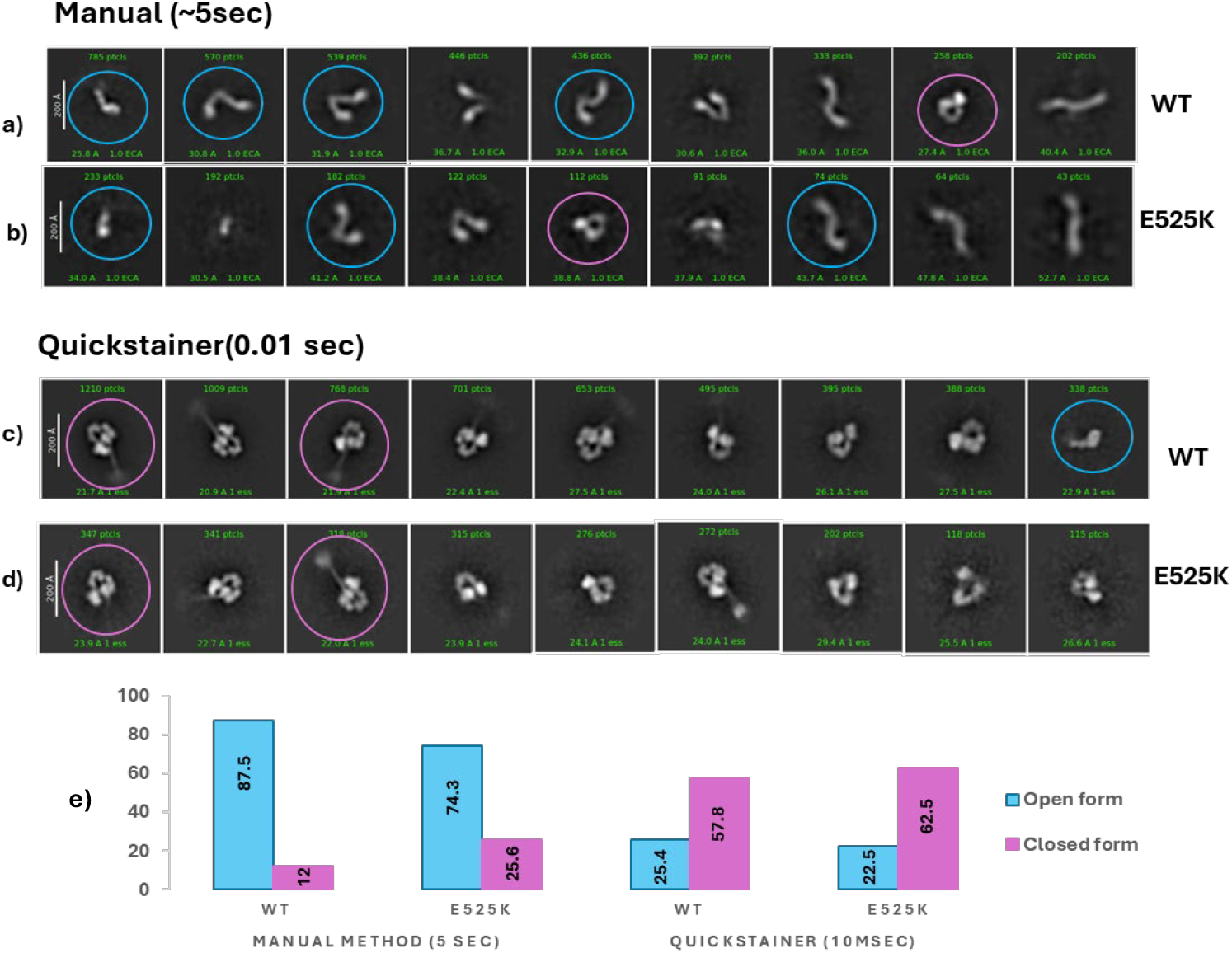
Comparing the effect of the myosin head variant E525K on IHM structure using manual and quick staining. **a, b**. 2D class averages of WT and mutant constructs using manual staining. Most molecules are open (blue) while a small number show interacting heads (closed, pink). The number of closed molecules approximately doubles with the variant (**e**, left). **c, d**. Same as **a, b**, only with QuickStainer. Closed structures are much more common overall (~60%), but with little difference between mutant and WT (**e**, right). **e**. Percentages of molecules in closed and open configurations using manual and quick staining. Similar, high numbers of WT and E525K molecules show the closed conformation with QuickStainer, consistent with FRET data under the same ionic conditions [10], demonstrating that short incubation times are better at preserving structure similar to that in solution; with manual staining, the IHM is mostly disrupted, especially in the case of the WT. These experiments were done with a 25-heptad myosin construct in contrast to the 15-heptad construct used in other figures and in [10].

Other approaches have also been used to address structural disruption caused by charged, carbon substrates grids. Crosslinking of the IHM in solution fully stabilizes the structure against disruptive forces [20, 28, 38] with minimal change in conformation [34, 36, 38]. However, crosslinking can artificially alter a closed-open equilibrium in favor of the closed conformer by stabilizing molecules when they enter the closed conformation (*cf*. [46]), which could lead to an inaccurate measure of the equilibrium constant. This problem is avoided by QuickStainer. The disruptive effects of a charged carbon substrate can also be avoided by observing protein specimens embedded in stain over holes in the carbon, where there is no interaction with substrate. We have demonstrated this with myosin filaments, which can become disordered upon incubation on carbon (**Fig. 8a, b**), but are well-preserved over holes [4, 44, 47, 48]. The disadvantage of this approach is that the stain stretching over the holes is unstable, and breaks in the electron beam, causing severe distortion. Coating with carbon after staining reduces this problem [4, 44, 47], but it can be avoided almost entirely with QuickStainer.

### Future Directions

There are several ways in which the QuickStainer can be further developed. The posts supporting sample and stain can be placed closer together (e.g., half the current distance), and a faster motor could be used, reducing incubation times below 10 ms, and enhancing the number of closed IHM structures (*cf*. **Fig. 4**). At the shortest times, the limiting factor may be the concentration of protein required to obtain an adequate distribution of particles. With further reduction in incubation time, it will be interesting to determine whether the fraction of closed structures reaches that predicted from solution studies [10]. Currently, it remains a puzzle why 40% of molecules are still open even with only 10 ms incubation: this suggests that opening of the closed structure is extremely rapid. QuickStainer could also be used to carry out time-resolved negative staining [9, 49, 50], for example capturing structural changes at different time points following binding of ligand to a protein molecule. In this case, the sample would be on the Protein post 1, ligand (e.g. ATP, Ca^2+^, etc.) on post 2, and stain on subsequent posts. Using the Arduino controller, dwell time could be adjusted on post 2 to obtain different ligand incubation times before staining. Finally, there are many examples of proteins, apart from myosin, that are also labile and are likely to be affected in similar ways using conventional (manual) negative staining. Quick staining should be useful in analyzing their structure and giving a more accurate idea of structural stability for cryo-EM optimization.

## ACKNOWLEDGMENTS

This work was presented in preliminary form [51]. We thank Drs Greg Hendricks and Kyoung Hwan Lee of the Electron Microscopy Facility at UMass Chan Medical School for help with microscope operation and training, and for providing the AAV sample. We are grateful to Drs Mitsuo Ikebe and Osamu Sato for smooth muscle myosin. This research was funded by the National Institutes of Health, grant numbers HL164560, AR081941, HL163585, and HL150953. The content is solely the responsibility of the authors and does not necessarily represent the official views of the National Institutes of Health.

## DATA AVAILABILITY

Plans for 3D printing of the QuickStainer parts are available at the Open Science Framework, https://osf.io/6d43b/?view_only=c55db1dd895e4785b813d80d5673aafe.

## AUTHOR CONTRIBUTIONS

Conceptualization: VN, RS, AS, DD, RP, RC; rotary stepper motor concept for quick staining, RP; design and construction of QuickStainer: VN; experiments using QuickStainer: RS, AS, AP, VN; analysis: RS, AS; protein preparation: JG, CMY; funding acquisition: RP, CMY; project administration: RC, RP; supervision: RC, RP, CMY; writing: RC, first draft; contributions from all authors to final draft.

## DECLARATION OF INTERESTS

The authors have no relevant financial or non-financial interests to disclose.

## Notes

### Competing Interest Statement

The authors have declared no competing interest.

